# Within-host adaptation of a foliar pathogen, *Xanthomonas*, on pepper in presence of quantitative resistance and ozone stress

**DOI:** 10.1101/2024.02.29.582737

**Authors:** Amanpreet Kaur, Ivory Russell, Ranlin Liu, Auston Holland, Rishi Bhandari, Neha Potnis

## Abstract

- The evolving threat of new pathogen variants in the face of global environmental changes poses a risk to the plant health and can impact the efficacy of resistance-based disease management.
- Here, we studied short-term eco-evolutionary response of the pathogen, *Xanthomonas perforans*, on quantitative resistant and susceptible pepper during a single growing season in open-top chambers under the influence of elevated Ozone (O3).
- We observed increased disease severity, accompanied by higher variation on resistant cultivar under elevated O3, with no apparent change on the susceptible cultivar. This altered resistance response under elevated O3 is linked to altered eco-evolutionary dynamics of pathogen. While a single pathogen genotype remained prevalent on susceptible cultivar, resistant cultivar supported heterogenous pathogen population, with the evidence of short-term evolutionary modifications seeded by *de novo* parallel mutations. Altered O3 levels led to strain turnover on resistant cultivar with higher within-host polymorphism containing higher proportion of random *de novo* mutations lacking parallelism.
- Population heterogeneity is a mechanism of pathogen adaptation in response to the stressors. While parallel mutations in response to quantitative resistance may provide clues to predicting long-term pathogen evolution, high proportion of transient mutations suggest less predictable pathogen evolution under climatic alterations.

## Introduction

Plants and pathogens constantly engage in a co-evolutionary arms race, where interactions of plant resistance (R) gene with corresponding pathogen avirulence (Avr) gene trigger the dynamics of avoidance detection and balancing fitness penalties facilitating long-term maintenance of polymorphism in Avr and R genes (Van der Plank, 1968; Leonard & Czochor, 1980; Vera Cruz *et al*., 2000; Leach *et al*., 2001; Mauricio *et al*., 2003; Bakker *et al*., 2006; Karasov *et al*., 2014). Plant resistance is a sustainable and cost-effective approach to manage pathogen outbreaks effectively in modern agricultural systems, however, the durability of the resistance depends on the nature of the resistance genes. Qualitative resistance breaks down rapidly within 3-5 years. Quantitative resistance, on the other hand, is deemed durable due to slower adaptation of pathogen to the polygenic traits encompassing additive multiple small-effect quantitative trait loci (QTLs) (McDonald & Linde 2002; Poland *et al.,* 2009; St. Clair, 2010; Brown & Rant, 2013). However, this process of pathogen adaptation to quantitative resistance is poorly understood (Pariaud *et al.,* 2009; Caffier *et al.,* 2016; Corwin & Kliebenstein, 2017).

Climatic fluctuations further add complexity to these eco-evolutionary dynamics of host-pathogen interactions and add uncertainty to the outcomes of resistance management. Climatic alterations can influence the stability of gene-for-gene interactions, can change the direction of selection (Cable *et al.,* 2017) with unexpected outcomes (Mostowy & Engelstädter, 2011). Global environmental changes can alter the evolution of virulence (Schmid-Hempel, 2003; Kutzer & Armitage, 2016), reinforce the selection on immunity genes, and accelerate pathogen adaptation thereby increasing plant disease risk (Laine, 2023). Climate-sensitivity of basal resistance (Janda *et al.,* 2019) as well as both qualitative and quantitative resistance (Webb *et al.,* 2010; Cheng *et al.,* 2013; Onaga *et al.,* 2017) has been noted, with few noted as climate-resilient (Cohen *et al.,* 2017). Climate sensitivity of plant pathogens within hosts is unclear (Huot *et al*., 2017; Onaga *et al.,* 2017; Velásquez *et al.,* 2018), and unlike animal pathogens (Shapiro & Cowen, 2012), little is known about virulence modulation under abiotic stress.

Field pathogenomics on spatio-temporal samples combined with population genetics has been the approach to study pathogen adaptation, although major focus has been the host selection pressure (Grünwald *et al.,* 2016; Badouin *et al.,* 2017; Hartmann & Croll 2017; Thilliez *et al*., 2019). These studies have informed that various mechanisms of genetic variation are in place, such as allelic polymorphisms in pathogen avirulence genes (Gassmann *et al.,* 2000; Huang *et al.,* 2014; Martynov *et al.,* 2019; Huang *et al*., 2019; Yang *et al.,* 2019; Gautier *et al*., 2023), genome rearrangements, with identification of rapidly evolving genomic regions carrying clusters of virulence determinants (Rouxel *et al.,* 2011; Whisson *et al.,* 2012; van Dam *et al*., 2016; Frantzeskakis *et al.,* 2020), gene gain/loss bringing in genetic novelty (Yoshida *et al*., 2016; Hartmann & Croll, 2017; Tsushima *et al.,* 2019), or sexual recombination facilitating rapid fixation of beneficial mutations (Grandaubert *et al.,* 2019). Distinct genomic compartments were found to have different rates and modes of evolution (Montoya & Raffaelli, 2010; Möller & Stukenbrock, 2017). Genomic signatures in the pathogen indicate influence of both directional and balancing selection in adaptation (de Vries *et al.,* 2020). Although characterized by transient polymorphism, directional selection leads to fixation of a single allele through selective sweep that selects for a higher fitness (Poppe *et al.,* 2015; Hall *et al*., 2020). Meanwhile, some virulence effectors are seen under diversifying or balancing selection maintaining alternative alleles within populations, leading to retention of sequence diversity (Brunner & McDonald, 2018). While the impact of host selection pressure in driving genomic changes has been explored, impact of global environmental changes driving pathogen evolution is limited (Wu *et al.,* 2020; Nnadi & Carter, 2021). Pathogens have an inherent ability to cope with rapid environmental changes, such as diurnal changes in temperature, UV, or water availability. Their response to these rapid fluctuations is considered plastic, for example, gene regulatory networks that allow growth and reproduction in response to fluctuating substrate availability. However, response to environmental changes occurring over longer timescales has led to specific microbial adaptations, resulting in increased phenotypic plasticity or genetic change. Seasonal turnover of specific pathogen genotypes with dominance depending on their fitness levels was demonstrated in the case of pseudomonads isolated from sugar beet leaves over the course of three growing seasons (Ellis *et al.,* 1999).

Most data on pathogen evolution comes from the temporal field surveys coupled with isolate genome sequencing, which may carry inherent culturing bias (Newberry *et al.,* 2020) and often do not dissect the impact of complex host, pathogen, and climate interactions. The strain-resolved metagenomics approach overcomes the bias of selecting only a dominant pathogen lineage and provides a comprehensive view of pathogen variants, including low-abundant variants, that may hold the potential to cause outbreaks under suitable environment (Newberry *et al.,* 2020). In this study, we used strain-resolved metagenomics to investigate within-host dynamics with high resolution into genetic processes shaping rapid pathogen evolution during a single growing season. We used an experimental setup in open-top chambers where we inoculated resistant and susceptible pepper cultivars with *Xanthomonas perforans* (*Xp,* also referred to as *X. euvesicatoria* pv. *perforans*) and exposed them to ambient and elevated ozone (O3) conditions (Fig. **1a,b**) and recorded disease severity levels twice during the season. As previously noted, elevated O3 did not influence disease severity levels on the susceptible cultivar (Bhandari *et al.,* 2023). However, under elevated O3 levels, the resistant cultivar showed significantly higher disease severity scores with an average increase of 12% during mid-season and 2% during end-season compared to an ambient environment. This increase was accompanied by high variation in disease severity, ranging from 2-38% during mid-season and from 0-8% during the end-season (Bhandari *et al.,* 2023) (Fig. **1c**). This high variation in disease severity levels could be a due to a plastic response of the resistant host under altered climate, direct or indirect effects of climate on pathogen infectivity or altered ecological interactions among pathogen genotypes, or combination of all these factors. In this study, we focused on investigating the influence of altered O3 levels on a pathogen population from a gene-to-population level and whether ecological fluctuations and/or evolutionary modifications in the pathogen population could explain the increase in disease severity observed on the resistant cultivar under elevated O3 conditions. The co-existence of multiple genotypes based on trade-offs or cross-feeding is proposed as one of the mechanisms for adapting bacteria to environmental stressors. We hypothesized that pathogen adaptation to the simultaneous challenge of host defense and elevated O3 would involve population heterogeneity and higher rate of evolution. To test this hypothesis, we chose to study coinfection using two closely related genotypes of *Xp*. The two selected genotypes were isolated during the recent field sampling (Newberry *et al*., 2023) and belong to two different sequence clusters (SCs) within the species, namely, SC6 (AL65, isolated from a susceptible pepper cultivar) and SC2 (AL22, isolated from a resistant pepper cultivar). The population genetic methods applied to the high-resolution shotgun metagenome data allowed us to capture the population dynamics at the intra-subspecific level of phylogenetic resolution, learn about the generation and maintenance of genetic variation and assess whether the variation could explain increased susceptibility on resistant cultivar under elevated O3. Overall, our data provide an understanding of how population dynamics both shape and are shaped by evolutionary processes.

**Figure 1.**
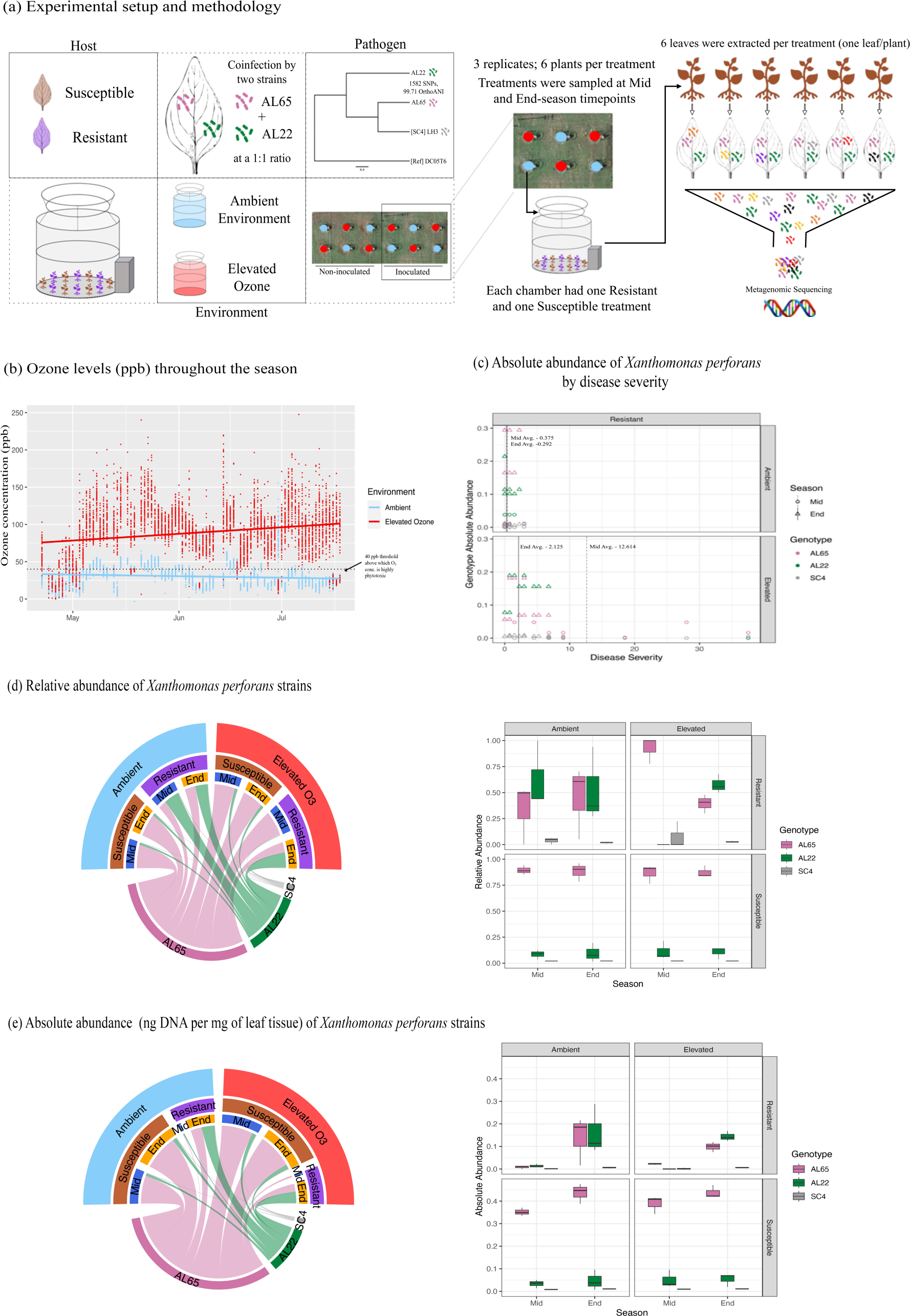
Experimental setup and pathogen population and disease dynamics. (**a**) We set up an experiment in which we inoculated two *Xanthomonas perforans (Xp) strains* at a 1:1 ratio onto susceptible and resistant plants and exposed them to ambient or elevated levels of O3 within open- top chambers. Out of 12 open-top chambers, six were non-inoculated controls, and six were inoculated (highlighted inside the black square box); additionally, six were exposed to elevated O3 (presented with red colored circles) while the remaining six were not exposed (shown with sky- blue colored circles). Each chamber contained 12 plants, making up two treatments, half susceptible cultivars and half resistant, with each cultivar set being a treatment, respectively. At mid and end-season time points, three samples per treatment were collected, with each sample comprising leaves picked randomly from six plants per chamber, and their metagenomic DNA was extracted and sequenced. The relative abundance of inoculated pathogen genotypes AL22 and AL65 was traced. We also observed evidence of natural infection by a representative of sequence cluster 4 (SC4), although it did not show an increase in abundance over time; (**b**) O3 levels throughout the season within the open-top chambers were on average 29.33 ppb (parts per billions) for Ambient chambers and 87.65 ppb for elevated O3 chambers. O3 levels above 40 ppb are considered to be highly phytotoxic (Saxena *et al*., 2020); (**c**) Absolute abundance of *Xp* genotypes as compared to disease severity during the mid and end-season sampling time points for the resistant cultivar under ambient versus elevated O3 levels. The mid and end-season disease severity averages vary from ambient to elevated O3; (**d**) Relative abundance of *Xp* genotypes as averages in the leftmost circular chord diagram and raw values in the rightmost barplot. (**e**) Absolute abundance (ng DNA per mg of leaf tissue) of *Xp* genotypes as averages in the leftmost circular chord diagram and as raw values in the rightmost barplot.

## Materials and Methods

### Data

The raw sequencing reads for metagenome samples used in this study were from Bhandari *et al.,* 2023 study (Accession number PRJNA889178), conducted in the 2021 growing season in open-top chambers at Auburn University. The detailed experimental design involving coinfection by two pathogen genotypes is described in Fig. **1a**.

### Abundance of inoculated strains

We assessed the influence of host genotype and elevated O3 on strain dynamics by tracing the relative and absolute abundance of two inoculated pathogen genotypes using StrainEst (Albanese & Donati, 2017). Briefly, the reference Single Nucleotide Variants (SNV) profiles representative of six lineages referred as sequence clusters (SC) within *Xp* (Newberry *et al.,* 2020). Representative strains chosen for each SC were GEV993 (SC1), AL22 (SC2), AL57 (SC3), LH3 (SC4), AL33 (SC5), and AL65 (SC6). An ordered vector of SNVs was obtained from SNV profile and SNV matrix of the species was constructed. This approach allowed the evaluation of natural infection, rather than limiting strain profiling of the inoculated strains. The metagenomic reads were aligned to extract the frequency of occurrences of each of the four possible alleles from the aligned reads, low coverage sites were removed, and the frequency profile for each metagenome was obtained using the lasso regression model (Albanese & Donati, 2017). This provided a relative abundance of different SCs in each sample. We calculated an estimate of absolute abundance of each SCs by multiplying their relative abundance with ng of DNA per mg of sample.

### Non-redundant pangenome of *Xp* inoculated strains

A non-redundant pangenome of *Xp* strains AL65 and AL22 was built using SuperPang (v0.9.4beta1) (Puente-Sánchez *et al*., 2022). This pangenome was then used as a reference for the rest of the analyses.

### Identification and removal of blacklisted genes from metagenomic reads

Since we were interested in inferring evolutionary changes in the coinfected *Xp* population, we implemented additional filters to avoid read stealing and donating. Related co-occurring *Xanthomonas* species or other phyllosphere microbes with more than 97% similarity in some housekeeping genes posed a challenge of read mis-mapping, hence an overestimation of parameters. Thus, we excluded these reads matching these high-similarity genes, referred to as blacklisted genes, from our analyses. To identify blacklisted genes, we executed the “run_midas.py species” command to identify the predominant species in the samples, utilizing the MIDAS (v1.3.2) default database as a reference (consisting of 31,007 bacterial reference genomes organized into 5,952 species groups) (Nayfach, 2022). Then, we built a custom database of pangenome using MIDAS and subjected it to BLAST (blast+) with the concatenated gene sequences from the species with a mean abundance > 0 (listed in Table **S1**) from the previous step.

The gene sequences with a similarity of 97% from the database were removed from all the samples using bbduk (v37.36) (Bushnell, 2014). The number of reads removed due to blacklisted genes are documented in the table (Table **S2**).

### Genetic differentiation and selection pressure

For following analyses, we mapped the samples to the pangenome using BWA-MEM (v0.7.12) (Li & Durbin, 2009), followed by the removal of low-quality alignments and duplicate reads using samtools (v1.11) (Li *et al.,* 2009) and picard (v1.79) (https://broadinstitute.github.io/picard/), respectively. The pairwise Fixation index (FST) was calculated for each 1kbp slide window using PoPoolation2 (v1201) with the parameters: --min-count 2, --min-coverage 4, and --max-coverage 120. To calculate within-host nucleotide diversity (π), Tajima’s D, and the ratio of non-synonymous to synonymous polymorphisms (pN/pS) for each gene, we used MetaPop (v1.0) (Gregory *et al.,* 2022). We employed MetaPop’s local alignment algorithm, which normalizes diversity estimates by dividing them by the genome length to account for uneven coverage across all samples. Also, it excludes SNV positions not covered in the genome length and sets a PHRED score threshold of > 20 for local SNV calls. For estimating population mutation rate per site (θ) (based on Watterson’s estimate), Rhometa (v1.0.2) was used with the reference database of pangenome. Rhometa uses dataset depth to calculate θ instead of the number of genomes as typically implemented in LDhat since an exact number of genomes is unknown in metagenome samples (Krishnan *et al.,* 2022). For statistics, the Shapiro test was run for the distribution of our dataset; the Kruskal-Wallis rank sum test and Dunn test were performed for overall and pairwise comparisons, respectively. A *P-value* of < 0.05 is denoted as statistical significance.

### Parallel evolution of SNVs differentiating across treatments and environments

Cochran–Mantel–Haenszel test statistics (CMH test) implemented in the PoPoolaton2 software was used to identify the consistent SNVs among all biological replicates with the parameters as mentioned earlier. SNVs with *P-values* above the Bonferroni-corrected significance threshold was considered outliers. Genome-wide SNVs were plotted in the Manhattan plot using ggplot2 (v3.4.2) in R.

### Evolutionary modifications

We calculated minor allele frequencies of SNV sites across the pangenome using MIDAS. First, we ran “midas.py snps” script for different samples and then used “merge_midas.py snps” to merge all replicates within treatments with the default settings to find major and minor alleles. Further, we removed the SNV sites associated with blacklisted genes and kept only those sites with a minimum read depth of 10 and present among at least two replicates. Mummer (v3.0) (Kurtz *et al.,* 2004) was used to identify genomes associated with different alleles within the population. For allele shift, sites, where the minor allele was shifted (frequency < 0.2) during mid-season to the major allele (frequency > 0.8) by the end-season were recorded. On the other hand, when major alleles recorded for mid and end-season did not match with alleles originating from AL22, AL65, and LH3, those were considered *de novo* mutations.

### Gene gain and loss

We used MIDAS (v1.3.2) with default settings to estimate gene changes in the pathogen population. A threshold of copy number of 0.35 was considered for determining gene presence (Garud and Pollard, 2020). Blacklisted genes were removed later from the output. All gene annotations were performed using eggNOG-mapper v2 (Cantalapiedra *et al.,* 2021).

## Results

### Pathogen population dynamics was host-genotype-dependent, and elevated O3 altered strain dynamics on the resistant cultivar

To understand pathogen population response during adaptation to the resistant host and under altered O3 levels, we traced the frequencies of the co-inoculated pathogen genotypes across treatments over time. Although there was an incidence of natural infection by the SC4 genotype, this genotype did not increase in frequency during the growing season. The presence of host resistance had a significant effect on the relative and absolute abundance of inoculated genotypes AL65 (*P* < 0.01, Kruskal-Wallis) and AL22 (*P <* 0.01, Kruskal-Wallis) (Fig. **1d,e**). Strain AL65 outperformed strain AL22 on the susceptible cultivar throughout the growing season, regardless of O3 level. On the other hand, strain-level heterogeneity was observed on the resistant cultivar, although the host x environment interaction likely influenced the strain dynamics. Strain AL22 (isolated initially from a resistant cultivar and thus can be assumed to be resistant cultivar adapted) maintained a higher population on the resistant cultivar under an ambient environment during mid-season. Despite the lower absolute abundance of *Xanthomonas* during the mid-season on the resistant cultivar, it is worth noting to observe the two strains with distinct niche preferences, with AL65 being dominant under elevated O3 and AL22 under ambient conditions. By the end-season, both strains co-existed on the resistant cultivar regardless of the environmental conditions (Fig. **1d,e**).

### Higher genetic differentiation in the pathogen population is observed on the resistant cultivar under elevated O3

Next, we looked whether genetic differentiation in the pathogen population may account for higher disease severity values seen under elevated O3 on the resistant cultivar since the absolute abundance of *Xanthomonas* could not explain the observation (Fig. **1c**, Bhandari *et al.,* 2023). Here, we hypothesized that adaptation to the stressors (host resistance or elevated O3 in either host background) is reflected in higher genetic differentiation in the pathogen population. From pairwise host comparisons, FST for pathogen populations recovered from susceptible cultivar was the lowest of all pairwise comparisons. Comparing pathogen populations from the resistant cultivar across O3 environments showed higher genetic divergence than that from the susceptible cultivar. Sampling time also influenced genetic differentiation levels, with mid-season populations having higher FST-values than end-season. However, adaptation to the resistant host under elevated O3 resulted in greater population divergence at the end-season. These observations for genetic differentiation were further confirmed with within-host nucleotide diversity values. We observed higher but variable within-host nucleotide diversity (π) values and mean population mutation rate per site (θ) in the pathogen population from the resistant cultivar as compared to the susceptible cultivar irrespective of environmental conditions (*P*π < 0.001, *P*θ < 0.001). End-season pathogen populations from the resistant cultivar under elevated O3 conditions had greater within-host nucleotide diversity values and mutation rates than mid-season populations (*P*π < 1e-12). Interestingly, nucleotide diversity for pathogen populations from the resistant cultivar during mid-season was significantly higher under ambient O3 conditions as compared to under elevated O3 (*P*π < 1e-5, *P*θ = ns), but there was no significant difference in the within-host nucleotide diversity values by the end-season (Fig. **2b,d**). The mutation rate on the susceptible cultivar, unaffected by O3 levels, averaged ∼0.87 x 10^-4^ by the end-season (mid-season ∼1.17 x 10^-4^). On the resistant cultivar, a higher mutation rate was observed throughout the growing season, ranging from ∼7.17 x 10^-4^ per site during mid-season to 1.98 x 10^-4^ per site by the end-season. Elevated O3 resulted in lower θ-value of 6.12 x 10^-4^ during mid-season but maintained higher levels (4.22 x 10^-4^ per site) by end-season on the resistant cultivar, compared to ambient conditions (Fig. **2e**). Next, we measured the extent of within-host polymorphism in the pathogen population by identifying alleles displaying intermediate SNV frequency (0.2-0.8) and classifying them as synonymous or non-synonymous mutations (Fig. **2f**). We compared normalized SNV counts (relative to the absolute abundance of *Xp* population) across treatments. Overall, a higher SNV counts were observed on the resistant cultivar compared to the susceptible cultivar under both environments (*P*(ambient): mid < 0.0001, end < 0.01; *P*(elevated O3): mid < 0.05, end < 0.001). There were no significant differences between non-synonymous and synonymous mutations across different conditions.

**Figure 2.**
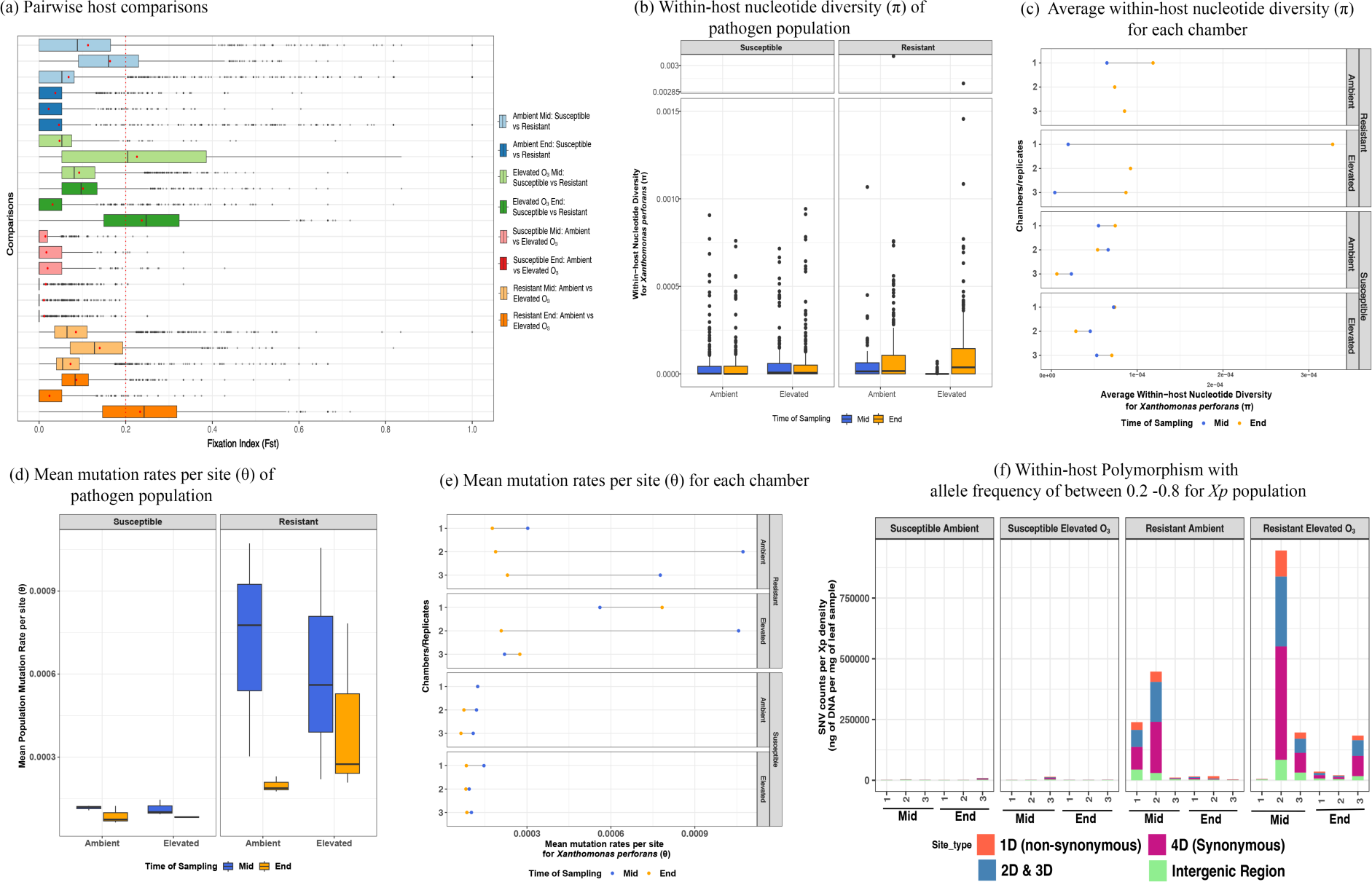
The pathogen population adapted to the resistant cultivar under elevated O3 with higher genetic differentiation. (**a**) Boxplots showing pairwise host comparisons with FST (Fixation Index) values (calculated using PoPoolation2) for 1kbp window. The red color dashed line indicates (values greater than 0.2) threshold value considered for differentiation; (**b**) Boxplot showing within-host nucleotide diversity (π) of *Xp* population together for different treatments; (**c**) Plot showing average nucleotide diversity (π) of *Xp* population for each chamber; (**d**) boxplots with mean mutation rates per site (θ) in *Xp* population; (**e**) Plot showing mean mutation rates per site (θ) in *Xp* population for each chamber; (**f**) Barplot indicating within host polymorphism with SNV counts (single nucleotide variant) for each chamber. These counts consist of different types of mutations (1D, 2D, 3D, and 4D) having an allele frequency between 0.2 and 0.8, where a site with 1D is one in which an amino acid change caused by nucleotide difference (non-synonymous), while a site with a 4D cannot be caused by any nucleotide difference (synonymous), and 2D & 3D indicates the either two or three possible changes, respectively can be tolerated, before an amino acid is altered (Chen & Garud, 2022; Nayfach, 2022).

### Signatures of parallel evolution in the pathogen population

The above observations of higher genetic differentiation and higher within-host nucleotide diversity values in the pathogen population recovered from resistant cultivar led us to examine further SNVs differentiating across host genotypes and O3 levels and evaluate the signatures of parallel evolution, i.e., SNVs observed independently across three replicates. More significant SNVs differentiating across host genotypes were identified compared to the number of SNVs differentiating across O3 levels with a Bonferroni-corrected threshold (Fig. **3a**). In the mid-season, we identified 171 significant SNVs (spanning 69 genes) and 109 SNVs (spanning 59 genes) that were distinctly observed when comparing pathogen populations from resistant and susceptible cultivars under ambient conditions and elevated O3 conditions respectively. By the end-season, these numbers dropped to just 29 (spanning 19 genes) under ambient conditions and 22 (spanning 15 genes) under elevated O3 conditions (Fig. **3b**). Among these 171 differentiated SNVs, only six SNVs (spanning six genes) were consistent across seasons, i.e., retained by the end-season under ambient conditions in genes encoding ribosomal protein RimK, molybdenum transport system permease protein, transketolase 1, a putative membrane protein, ribonuclease T, and Molybdenum cofactor guanylyltransferase. Under elevated O3 conditions, two SNVs differentiating across the pathogen population from the two host genotypes were observed to span gene encoding molybdenum cofactor guanylyltransferase and an intergenic region upstream of platelet-activating factor acetylhydrolase IB subunit genes (Fig. **3c,d**). Surprisingly, there were no SNVs under parallel evolution retained throughout the season, that were differentiating across resistant ambient and resistant elevated O3 conditions.

**Figure 3.**
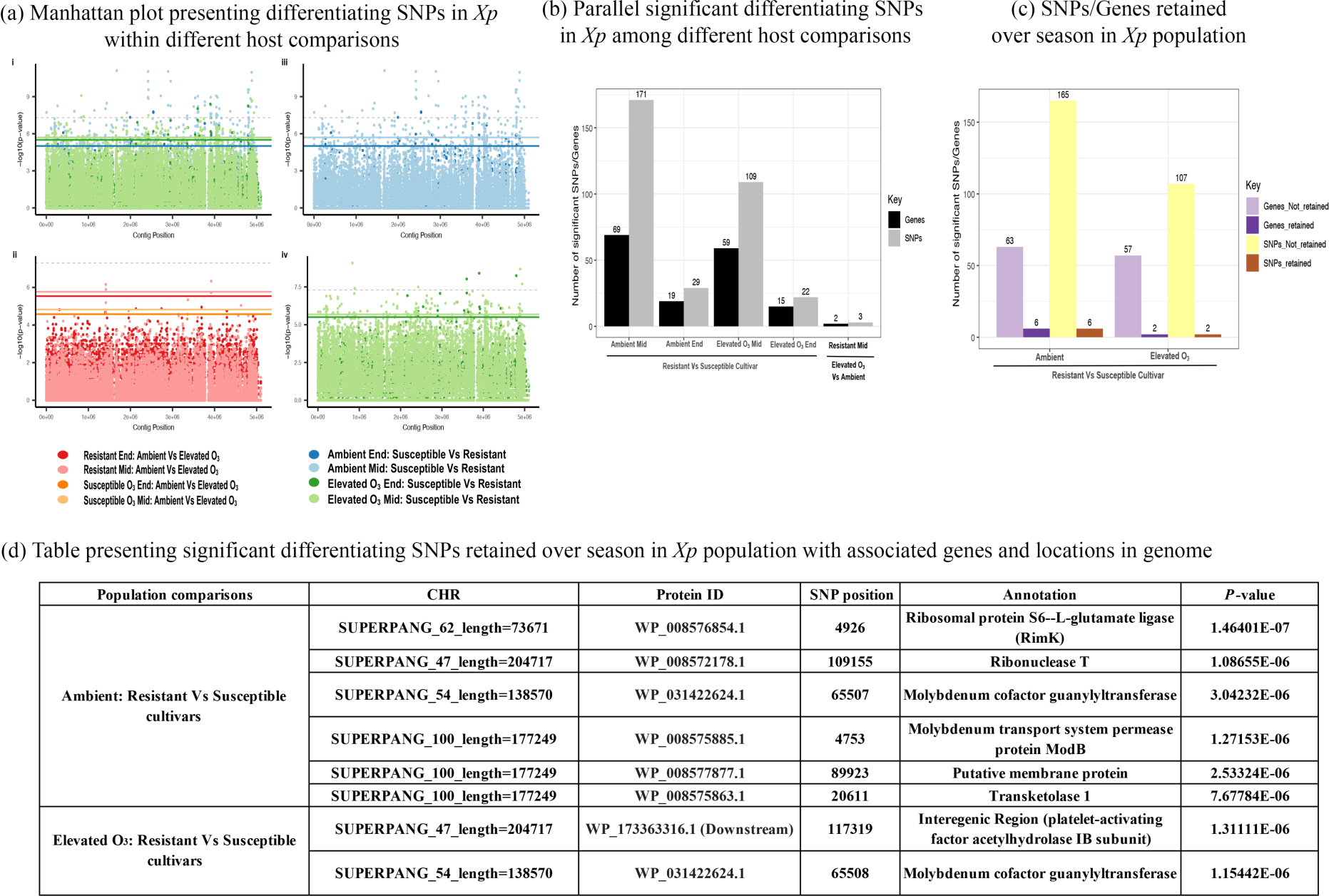
**Differentiating SNVs in pathogen populations were retained over season comparing host genotypes**. (**a**) Manhattan plot indicates differentiating SNVs across different host genotypes and environments. The grey dashed line is the default 5 x 10^-8^ GWAS threshold. The colored lines present the Bonferroni-corrected threshold, which is 0.05 divided by the number of SNVs in the summary statistics. (**b**) Barplot indicates the total number of SNVs and their associated genes were parallel across different comparisons. (**c**) Barplot showing the total number of SNVs and their associated gene counts persisted by the end-season or counts which revert to the original state or lost from pathogen population by end-season; (**d**) Table presenting all the SNVs associated genes with annotations and locations in the genome which were persistent across season.

### A strong purifying selection in response to altered environment

We delved deeper to elucidate the mechanisms involved in increased genetic differentiation observed within the *Xp* population. Our hypothesis centered on the possibility of parallel positive selection occurring within the pathogen population in response to both stressors. To test this hypothesis, we analyzed selective pressures within the genomic landscape, considering metrics such as Tajima’s D and pN/pS (non-synonymous to synonymous) values to uncover their contributions to the observed genetic differentiation. We detected significant differences in pN/pS ratios based on cultivar (*P <* 1e-5), season (*P <* 1e-5), and environment conditions (*P <* 1e-8). We observed that pN/pS ratios in populations from resistant cultivars exposed to elevated O3 conditions differed significantly from those in resistant cultivars under ambient conditions (*P <* 1e-4) and susceptible cultivars under elevated O3 conditions (*P <* 0.01). Then, we categorized the genes according to their selection pressures, distinguishing between those under positive (pN/pS >1), negative (pN/pS = 0), and purifying selection (pN/pS < 1). Metapop used a coverage threshold of less than 70% of their length covered and less than 10× average read depth coverage, which led to the removal of samples from resistant cultivars (2 replicates of mid-season under both conditions and 1 replicate of end-season under elevated O3). Approximately ∼98.85 % (average of 4096 genes, range 4079-4520 genes) of the pangenome (4603 genes) showed finite and defined pN/pS ratio values across all the samples. Of these, ∼ 99.8% of genes exhibited purifying (stabilizing) selection (pN/pS < 1) regardless host and environmental conditions. Within purifying selection, about 98% of genes exhibited strong negative selective pressure (pN/pS = 0) in the pathogen genome, indicating the pathogen population maintained strong evolutionary conservation by maintaining their gene functions and natural selection effectively filtered out deleterious or harmful mutations within the pathogen population (Fig. **4a**). The proportion of genome under positive selection (pN/pS > 1) was relatively small with an average of 0.2% genes (ranging from 2-13 genes) in pathogen population from susceptible cultivar irrespective of conditions. Similar numbers were observed for pathogen from resistant cultivar (1-19 genes). Tajima’s D values were significantly different in the resistant host under elevated O3 conditions compared to the resistant cultivar under ambient conditions (*P <* 0.05), as well as the susceptible cultivar under both ambient (*P <* 0.05) and elevated O3 conditions (*P <* 1e-15). The highly negative values of Tajima’s D in resistant cultivars by the end-season, regardless of conditions, indicate the presence of a selective sweep or rare alleles in the population (Fig. **4b**). Next, we scanned for genes experiencing positive selection (with pN/pS values >1 and Tajima’s D < 0) across independent replicates and retained from mid to end-season. Low sequencing depth for some of the mid-season samples from resistant cultivar did not allow us to monitor parallel evolution on resistant cultivar. However, parallel evolution was observed in 12 genes in the pathogen population from susceptible cultivar under elevated conditions and two under ambient conditions (Fig. **4c**). Of those, acid phosphatase and von Willebrand factor type A were under parallel evolution under ambient and elevated conditions. Genes identified under parallel evolution under elevated O3 conditions on the susceptible cultivar included transcription regulators, sigma factor, SAM-binding methyltransferase, aminopeptidase, and transporters.

**Figure 4.**
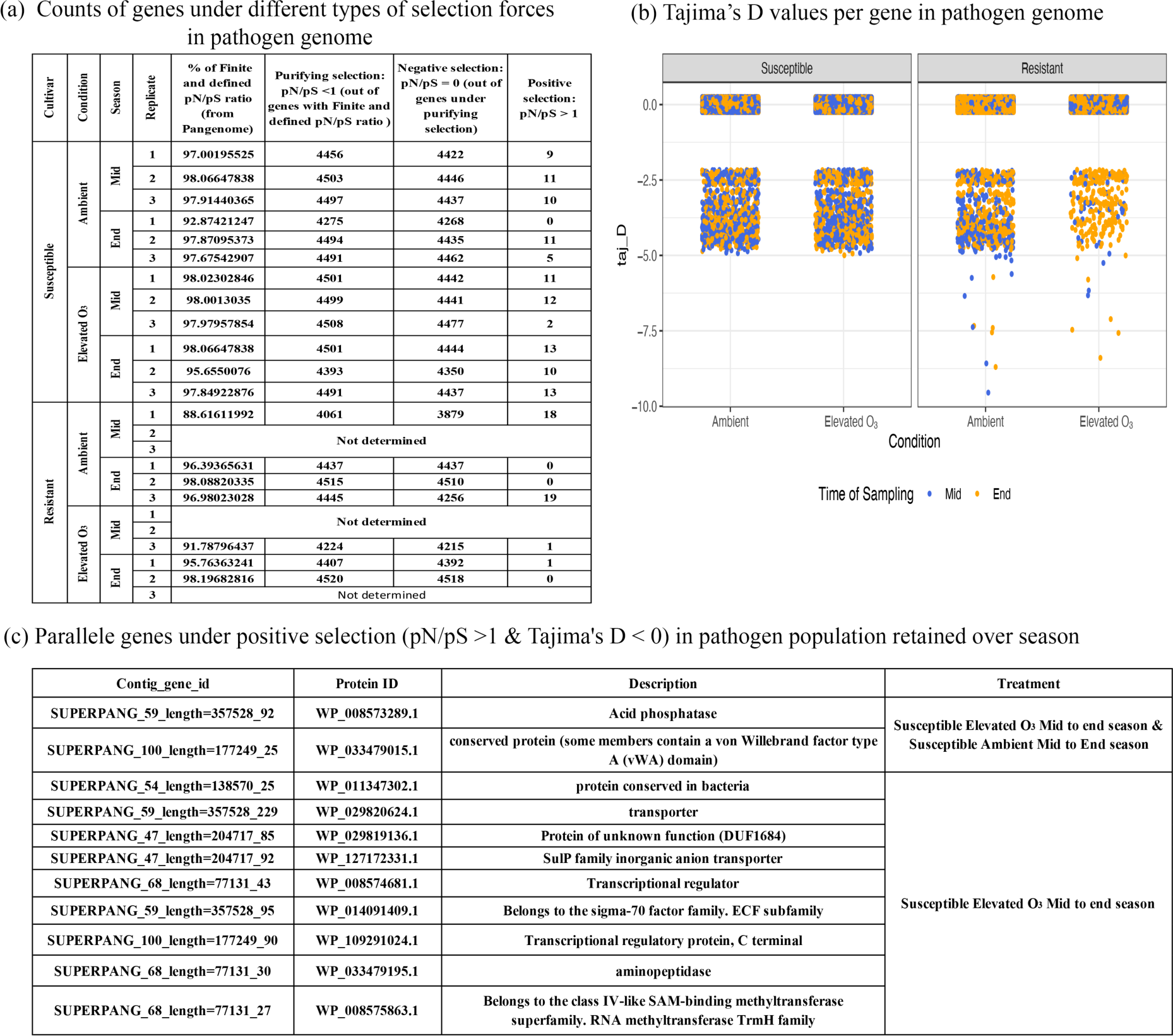
Evidence for purifying selection. (**a**) Table presenting counts of genes in *Xp* population under different selection types: negative selection or strong purifying selection (pN/pS = 0), positive selection (pN/pS >1) and purifying selection (pN/pS between 0 to 1); (**b**) Point graph presenting the Tajima’s D values per gene across different treatments; **(c)** list of genes with their NCBI protein Ids under positive selection (pN/pS >1 and Tajima’s D <0) in pathogen population across different treatments.

### Genetic differentiation observed across cultivars and O3 conditions is due to standing genetic variation and *de novo* mutations

The higher proportion of within-host polymorphism observed on the resistant cultivar (Fig. **2f**) may be attributable to standing genetic variation and/or *de novo* mutations. Indeed, results on strain dynamics indicated the co-existence of two pathogen genotypes on resistant cultivar and shifts in their frequencies under elevated O3, thus contributing to standing genetic variation (Fig. **1**). However, the contribution of evolutionary modifications that arose *de novo* during a growing season cannot be ruled out. Our experimental design involving co-inoculating two closely related pathogen genotypes (1582 SNVs apart) allowed us to assign the alleles to the parent genomes and tease apart SNVs arising during the growing season. To understand the contribution of allelic shifts that result from standing genetic variation, we focused on identifying SNVs with a frequency of ≤ 0.2 during mid-season but shifted to ≥ 0.8 by the end-season. These rare sweeping alleles may have adaptive potential. These alleles matched the AL65 and AL22 parent genomes, thus indicating the contribution of standing genetic variation. As expected from strain dynamics data, no such alleles were found in the susceptible cultivar, regardless of environmental conditions. However, a larger proportion of parallel allele shifts leading to fixation of AL22 alleles (134 alleles) was observed on resistant cultivar under elevated O3. These were associated with genes involved in host immune suppression or enhanced nutrient acquisition or assimilation, such as TonB-dependent receptor, outer membrane protein, XopV effector protein, and acid phosphatase (Jungnitz *et al.,* 1998; Deb *et al.,* 2020; Bhadouria & Giri, 2022). In contrast, the contribution of allele shift leading to fixation (4 alleles) was much smaller in pathogen on the resistant cultivar under ambient environment, with two alleles belonging to AL65 in the gene encoding XopAD type III effector protein (Table **S3**).

To identify if any SNVs resulted from *de novo* mutations, we identified sites with alleles that did not match any AL65 and AL22 parents and LH3 genome (natural infection). Of these *de novo* mutations, we first focused on those observed as novel major alleles in the end-season samples, parallel in at least two replicates with a minimum depth of 10. Due to variable sequencing depths across metagenome data from resistant and susceptible cultivars, we normalized the SNV counts with the average mean coverage/sequencing depth value per treatment. Normalized parallel de novo mutations were highest in the pathogen population recovered from resistant cultivar under ambient conditions (Fig. **5c**). Interestingly, normalized parallel mutation counts from resistant cultivar under elevated O3 were comparable to that from susceptible cultivar under ambient and elevated O3. We classified *de novo* parallel mutations into (1) those retained from mid to end-season, indicating their selection throughout the growing season, and (2) those that appeared after mid-season. Due to low sequencing depth, some *de novo* parallel mutations observed in population from resistant cultivar could not be assigned to either of these categories. The sites retained over time were then compared for their presence across host genotypes and O3 conditions, both in terms of their site ids and the associated gene id spanning the mutation sites. We identified 25 sites that spanned 12 genes that showed the presence of parallel *de novo* mutation in all treatments regardless of host genotype and O3 conditions. These sites were predominantly associated with signaling pathways known to play a crucial role in virulence, aiding pathogens in promoting disease within the host (Li et al. 2019; Wang & Qian 2019). There were 72 sites spanning 35 genes carrying *de novo* mutation in pathogen population from susceptible cultivar common in both environments. These included genes related to overall pathogen fitness, including DNA repair photolyase, outer membrane protein beta-barrel family, multidrug efflux RND transporter permease subunit, and peptidoglycan biosynthesis. Six sites spanning one gene and two intergenic regions were uniquely identified as selected sites in the population recovered from resistant cultivar under ambient environment. These were in genes encoding chemotaxis protein and monovalent cation proton antiporter 2 (CPA2) transporter (TC 2.A.37) family. Three *de novo* mutation sites retained from mid to end-season on resistant cultivar under elevated conditions were also found on susceptible cultivar in both environments (Fig. **5e**). These sites were found in the intergenic region of two genes coding for cache2/3 fusion domain-containing protein, which may assist the pathogen in locating nutrients under epiphytic conditions on plant leaves and identifying potential entry points into host cells (Brewster *et al.,* 2016) and encoding dienelactone hydrolase involved in chlorocatechol degradation (Schlömann *et al*., 1993). These are of interest given the observation of higher disease severity on resistant cultivar under elevated O3. There were parallel *de novo* mutations under elevated O3, linked to genes such as sulfotransferase family, peroxiredoxin activity, and carboxymuconolactone decarboxylase family protein, known for an antioxidative response in the host (Mougous *et al.,* 2002; Chen *et al.,* 2015; Kamariah *et al.,* 2018). We found one unique mutation in the pathogen population from a resistant cultivar under elevated O3 by the end-season, which was related to an intergenic region of genes known as TldD protein, identified under different stresses (Vobruba *et al.,* 2022). In addition, possible deleterious sites were identified based on their appearance in mid-season but lost by the end-season. These putative deleterious mutations were observed at higher normalized counts in populations under elevated O3 regardless of host genotype (Fig. **5c**, Table **S4, S5**).

**Figure 5.**
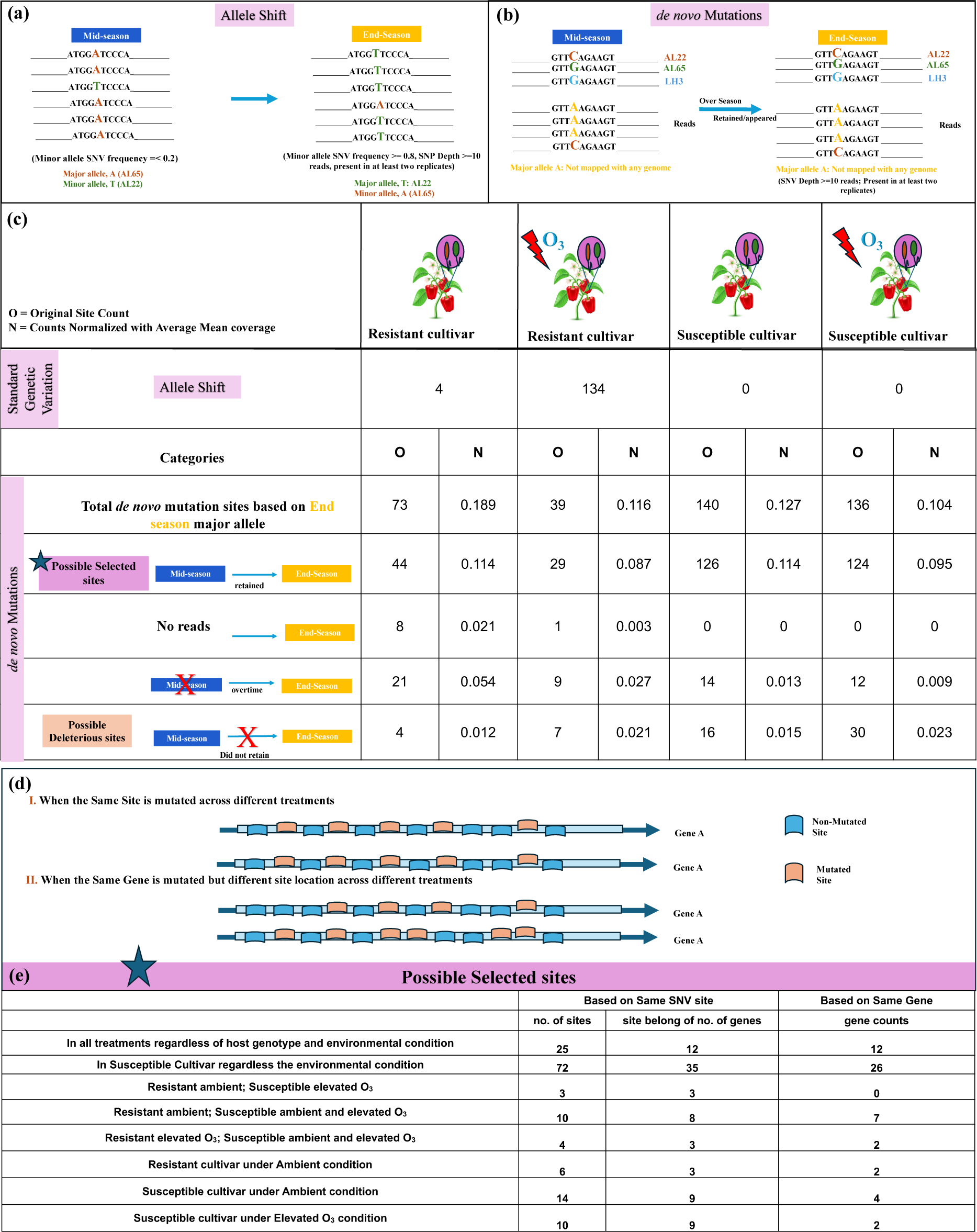
Evolutionary modifications with strain replacement exhibited in pathogen population in resistant hosts under elevated O3. (a) Visual representation of the allele shift in the *Xanthomonas perforans (Xp)* population, where a minor allele belonging to a specific site had a 0.2 as maximum frequency during the mid of the season, meaning it was present 20% or less number of population reads. By the end-season, however, the minor allele had transformed into a major allele with a frequency of at least 0.8, meaning it was present in 80% or more reads in population; **(b)** Visual representation of *de novo* mutations, in which the major alleles for mid and end-season were mapped to the reference genomes of *Xp* (AL22, AL65, and LH3) for all sites, and those that did not match any of the genomes were then pooled out; **(c)** Table presenting counts for the allele shift, and total *de novo* mutations for different treatments, which were further separated into different categories: (1) total *de novo* mutations were counted based on the mapping of major alleles during end-season, (2) out of total how many mutations retained in the population from mid to end-season (possible selected sites), (3) where we did not have mid-season information, (4) mutations only appeared over time by end-season, (4) possible deleterious allele sites, which did not retain by end-season in different treatments. We normalized the counts by dividing them by the average mean coverage/sequencing depth value per treatment. Note: In all the count estimations, only those sites with SNV depth of a minimum of 10 reads during the end-season with their presence in at least two replicates for all treatments were considered. Also, sites belonging to blacklisted genes were removed from the calculations; **(d)** Visual graphics for comparisons when the same site hit in parallel across different treatments; when the same gene was mutated across different treatments irrespective of site position; **(e)** Table at the bottom includes counts related to the sites that retained from mid-season to the end-season in population. These counts are based on when parallel *de novo* mutation appeared across different treatments based on the same position site and when the same gene belonging to parallel *de novo* mutations was mutated across different treatments.

### A higher rate of gene flux events was observed in the pathogen population during adaptation to resistant cultivar

Next, we measured changes in the gene pool, indicating variations in pangenome size. Our hypothesis posited that selection pressure would induce alterations in the pathogen gene pool, favoring gene acquisition or loss for improved pathogen fitness, thus we anticipated more gene gain/loss events in pathogen populations under both stressors. We defined “gene gain” as a gene reappearing from mid-to-end-season and “gene loss” as a gene present in mid-season but lost by end-season. We observed a higher gene flux in the pathogen population from resistant cultivars, especially under elevated O3 conditions. However, there were no differences in gain/loss events in pathogen population from susceptible cultivars irrespective of environmental conditions (Fig. **6a**). We identified a total of 90 gene gain and 81 gene loss events in resistant cultivar under elevated O3, out of which primarily gene gain events occurred in AL22, and losses were in the AL65 genome (Fig. **6b**). We found evidence of parallel gene flux events i.e observed across all three replicates. Around 20 core genes were lost in pathogen population from resistant cultivar under elevated O3. Around 8-10 core genes were lost in pathogen population from susceptible cultivar from either environment. Interestingly, genes belonging to COG categories M, GM, N, U, and P were lost from the susceptible cultivar, which were related to metabolism, transport, cell wall, cell motility, and intracellular trafficking. In contrast, M and U were gained in resistant cultivar under elevated O3. Additionally, genes belonging to C, P, K, and L categories were gained in the pathogen population from resistant cultivars under elevated O3 conditions with the adaptive advantage of energy production, transcription, nucleotide metabolism, replication, and repair (Fig. **6c**). Although seen as a lack of parallel genes being lost from pathogen population from resistant cultivar under ambient conditions, this may have been artifact due to low sequencing depth for some resistant cultivar samples (Fig. **6b**).

**Figure 6.**
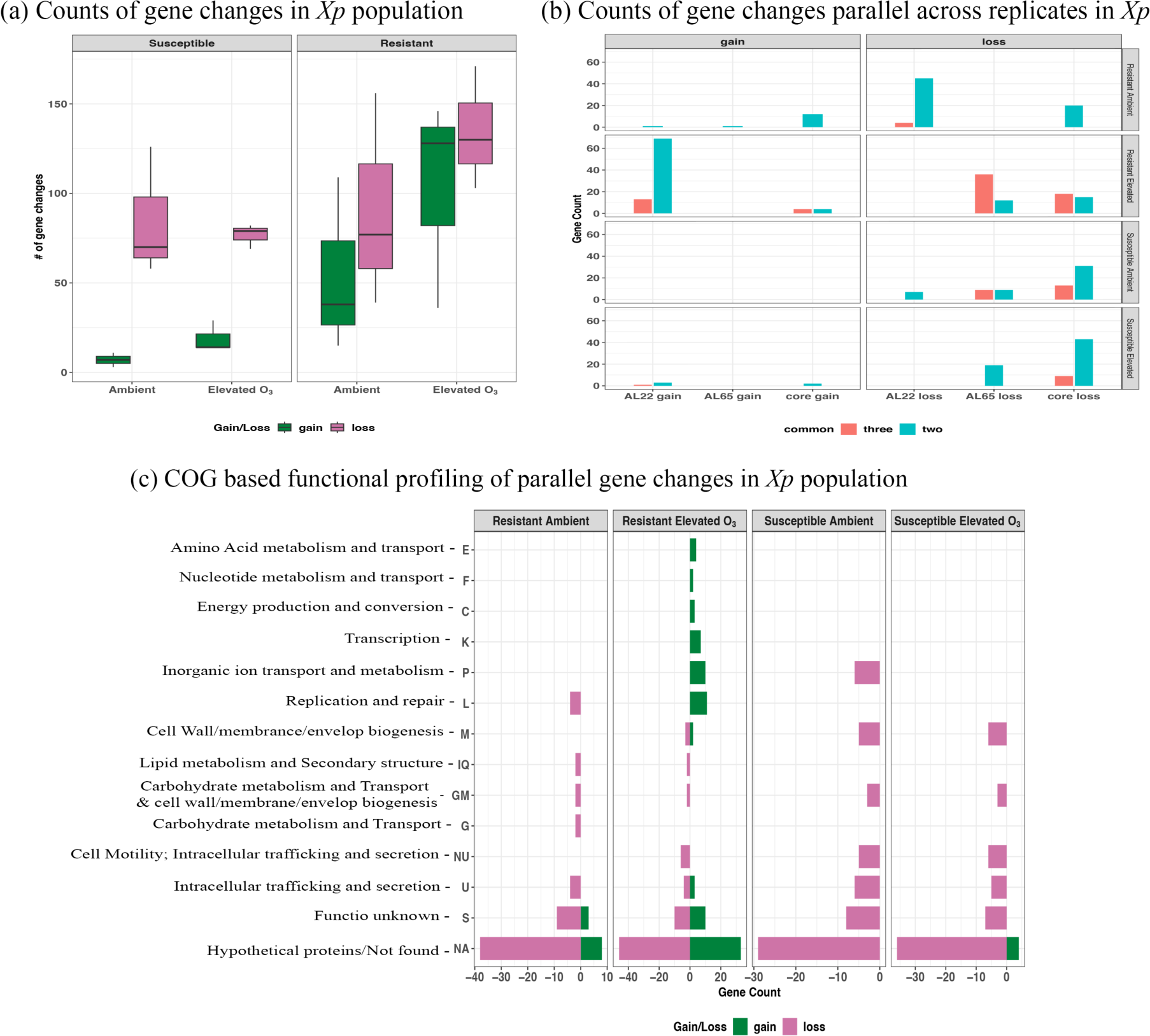
Gene flux in the pathogen population. (**a**) Boxplot showing the number of gene gains (absent during mid-season and present by end-season in the pathogen population) and gene losses (present during mid-season and lost by the end-season) across all the chambers. (**b**) Bar plot presenting a number of gene changes common among two and three chambers/replicates of different treatments. (**c**) Bar plot showing the COG (Cluster of Orthologous Groups) based functional profiling of gene changes common in two and three chambers across different treatments.

## Discussion

Global climate change can affect infectious diseases in a nonlinear fashion. However, we have little evidence that climate change has already led to a surge of infectious diseases as the globe has warmed up over the past century. Models have contrasting predictions of increases in the geographic range of infectious diseases, range shifts in disease distributions (Lafferty, 2009), or overall increased pathogen virulence (Velásquez *et al.,* 2018). Rapid and gradual changes in the environment can affect multiple scales of plant-pathogen interactions from the molecular to the community level, with feedback linking across these scales and adding uncertainty to the outcomes of resistance management. In this study, we harnessed the power of metagenomics to monitor the generation and maintenance of genetic variation in response to altered climate (elevated O3) and host selection pressure, its impact on phenotypic changes, and the fitness on which natural selection acts. The experimental setup allowed us to gather empirical evidence on rapid evolution in a single growing season under future climatic conditions in the open top-chamber field setup.

Elevated O3 did not influence disease severity on susceptible cultivar and did not alter pathogen population structure. A single pathogen genotype, AL65, dominated the susceptible cultivar throughout the growing season. While resistant cultivar supported the co-existence of both pathogen genotypes throughout an ambient environment, the strain turnover was observed under elevated O3, with AL65 being dominant during mid-season, followed by the co-existence of both strains by the end-season. The colonization success of AL65 on the resistant cultivar during mid-season was linked to higher disease severity under elevated O3, although accompanied by high variation (Fig. **1**). The altered pathogen dynamics on the resistant cultivar under elevated O3 could be a result of either altered host selection pressure or altered ecological interactions among the two closely related strains or altered interaction of individual strains with the host with changing host defense response under abiotic stress (Castelijn *et al*., 2012; Papkou *et al.,* 2021; Yang *et al*., 2022). Interestingly, despite the end-season pathogen population being heterogenous in both environments, phenotypic differences were evident in the two environments, with resistant plants under elevated O3 showing a 2% increase in disease severity compared to ambient conditions. While the absolute abundance of *Xanthomonas* could not solely explain this difference (Bhandari *et al.,* 2023), we investigated whether the amount of genetic variation available for selection to act on differed across environments and whether adaptation to the new environment was driven by new alleles of large effect that could be linked to the phenotype. Presence of mixed pathogen population and maintenance of heterogeneity in response to stressors has been referred to as a mechanism of rapid evolution resulting in higher pathogen fitness (Hiramatsu *et al.,* 2001; Balaban *et al.,* 2004; Aertsen & Michiels 2005; Longo & Hasty, 2006; Dhar & McKinney, 2007; Lidstrom & Konopka, 2010; Schröter & Dersch, 2019; Caballero-Huertas *et al.,* 2023). This co-existence may suggest that the interplay of two pathogen genotypes may contribute to the adaptation to small-effect quantitative trait loci on the resistant cultivar.

We find that elevated O3 alone was not the primary driver of genetic differentiation in the pathogen population, as suggested by stable mutation rate and within-host nucleotide diversity of *Xp* on susceptible cultivars under both environments. However, host selection pressure on the resistant cultivar led to relatively higher nucleotide diversity and a higher population mutation rate. The elevated O3 did not affect the average level of polymorphism, although the end-season population displayed a higher mutation rate compared to ambient environment. Elevated O3 also supported high variation in levels of polymorphism and mutation rates on resistant cultivar, signifying genotypic plasticity (Fig. **2**). This plasticity may suggest adaptive strategy in presence of stressors, as observed in various biological systems such as intestinal microbiota, coral reefs, marine life, and wildlife populations (Candela *et al.,* 2012; Reusch, 2014; Torda *et al.,* 2017; Leray *et al.,* 2021; Bonachela *et al.,* 2022; Risely *et al.,* 2023;) Next, we identified niche differentiating parallel SNVs retained throughout the growing season. Putative adaptive SNVs differentiating between resistant and susceptible cultivar under ambient environment were of interest. These SNVs-spanning genes encoded transcriptional factor, RimK, capable of converting environmental signals into dynamics changes in translational output and adaptive remodeling of proteome through specific modification of bacterial ribosomes (Little *et al.,* 2016), and molybdenum cofactor guanylyltransferase, involved in conversion of substrates generated in the host during inflammation, in response to defense-activated ROS-mediated defenses (Zhong *et al.,* 2020). These and other differentiating SNVs may have a role in pathogen adaptation to quantitative resistance. However, no such adaptive differentiating SNVs were retained over the season when comparing pathogen population across ambient and elevated O3. This was because, despite a greater level of polymorphism, many of these sites were inconsistent across replicates and many SNVs were lost throughout the growing season. A strong purifying selection in the pathogen population was observed irrespective of cultivar and conditions, acting against newly emerged harmful mutations, preserving the genetic traits, and leaving imprints on genetic diversity by alteration in the distribution of genetic variants at specific sites (Fig. **3**) (Cvijović *et al.,* 2018). While extremely negative Tajima’s D values suggested positive selection, low sequencing depth for some samples was a shortcoming preventing the identification of signatures of parallel evolution in pathogen from resistant cultivar. The presence of elevated O3 led to evidence of positive selection in pathogen population on susceptible cultivar in genes encoding transcriptional regulators, sigma factors, and TrmH (methyltransferase involved in post-translational regulation influencing oxidative stress response, and suppression of host immune response (Fig. **4**) (Rimbach *et al.,* 2015; Galvanin *et al.,* 2020). The evidence of selection was not limited to the nucleotide level, but selection at the gene level with parallel gene gain/loss events was evident. Variations in the rate of gene gain and loss have impacts on the pathogen’s fitness (Brockhurst et al. 2019; Domingo-Sananes and McInerney 2021; Lefébure and Stanhope 2007; Moulana et al. 2020). Pathogen adopted a strategy of “less is more” when adapting to resistant cultivar and under elevated O3 conditions that led to more gene loss events (Fig. **6**) (Li *et al.,* 2017; Seidl & Thomma, 2017; Simonsen, 2022). This observation of the co-existence of both pathogen genotypes and more gene loss events on resistant cultivar also aligns with the black queen hypothesis that states that species turnover and interactions among members in a diverse community select for loss of genes redundant in function and promote interdependence (Morris *et al.,* 2012).

While we observed evidence of the high level of polymorphism and signatures of positive selection in response to elevated O3 or host selection pressure, we categorized the observed genetic variations into those arising from standing genetic variation (those observed due to ecological changes resulting in strain-specific allele shifts) and from evolutionary modifications via retention of parallel *de novo* mutations throughout the growing season. Here, we focus on short-term single-season within-host changes in pathogen populations, which may provide insights into pathogen adaptation over longer time scales. The fact that we observe evidence of parallel evolution with allele shifts or *de novo* mutations that are retained over the growing season across replicates suggests that short-term evolutionary forces could contribute to signals of adaptation that accumulate over the long term. For example, parallel evolutionary signatures in genes specifically implicated in functions to overcome host defense, ROS, or enhanced nutrient assimilation might contribute to continuum responsible for resistance erosion in the longer term. We find that both standing genetic variation and *de novo* mutations play a role in rapid adaptation of pathogen onto quantitative resistant cultivar, although the contribution of each of these mechanisms of variation was variable depending on O3 environment. As strain turnover was observed under elevated O3 conditions, a larger contribution of standing genetic variation was evident (Fig. **5**). Examples of standing genetic variation in response to stressors contributing to rapid evolution through small allele frequency changes at multiple loci are common across prokaryotes or eukaryotes (Dayan *et al.,* 2019; Chen & Garud, 2022). We find that adaptation to resistant cultivar under ambient environment involved signatures of parallel *de novo* mutations, many of which are retained throughout the growing season. A reverse-genetics approach to test for the fitness effects of these retained mutations on resistant cultivar may help understand the adaptation process to quantitative resistance. Interestingly, although we observed a greater proportion of SNVs on resistant cultivar under elevated O3, some *de novo* mutations did not persist throughout the growing season, suggesting possible deleterious mutations, or lacked signatures of parallel evolution across replicates. These higher proportion of transient mutations may explain high degree of variation observed in disease severity levels across replicates under elevated O3. These findings suggest that abiotic stressors may have influence in altering pathogen evolution with unpredictable outcomes. Multi-seasonal experiments under OTC conditions by subjecting pathogen to altered climatic conditions are needed to understand the fate of these transient mutations in terms of their fixation and whether these small effect loci contribute towards further erosion of quantitative resistance.

In summary, patterns of genetic variation observed in this study may help in predicting the evolution of the pathogen and predicting the durability of disease resistance. However, the finding of random transient mutations leading to higher within-host polymorphism may point to difficulties in predicting pathogen evolution under climatic shifts.

## Acknowledgments

We thank the Alabama Agricultural Experiment Station for the experimental site and the hatch program of the National Institute of Food and Agriculture, United States Department of Agriculture for funding. We thank members of the Potnis lab for their help with sampling. This work was made possible in part by a grant of high-performance computing resources and technical support from the Alabama Supercomputer Authority.

## Competing Interests

The authors declare no competing interests.

## Author Contributions

NP conceptualized and designed the study. AH and RB contributed to the experimental design, conducted sample collections and processing of samples. AK and NP conceptualized the specific analyses presented in this study. AK, IR, RL conducted analyses, specifically, IR conducted ecological dynamics analysis. AK and IR worked on building pangenome and removal of blacklisted genes. RL conducted the PoPoolation analysis for SNV differentiation. AK conducted the selection analysis, gene gain/loss, polymorphism extent, and evolutionary modification. AK, IR, and NP worked on data interpretation. AK and NP wrote the manuscript with contribution from all the authors.

## Data Availability

Sequence data generated from this work have been deposited in the SRA (Sequencing Read Achieve) database under the BioProject accession PRJNA889178. *Xp* strain AL22 has been submitted under BioProject accession PRJNA1077988. *Xp* AL65 used from submission under BioProject accession PRJNA953417. All other data and code used in this study are available in the following GitHub repository (https://github.com/Potnislab/pathevo)

## Supporting Information

Table S1. Microbial species which had more than 0 mean abundance within the samples.

Table S2. Reads counts before and after removing blacklisted genes from samples.

Table S3. Parallel sites (across replicates) belonging to Allele-shift (with < 0.2 allele frequency during mid-season and became > 0.8 by the end-season)

Table S4. Parallel de novo mutation sites appeared across different conditions at the same position in the pathogen population genome.

Table S5. Genes belonging to Parallel de novo mutations: when the same gene was mutated across different treatments in the pathogen population genome.

## Notes

### Competing Interest Statement

The authors have declared no competing interest.

